# Species dynamics and interactions via metabolically informed consumer-resource models

**DOI:** 10.1101/518449

**Authors:** Mario E. Muscarella, James P. O’Dwyer

## Abstract

Quantifying the strength, sign, and origin of species interactions, along with their dependence on environmental context, is at the heart of prediction and understanding in ecological communities. Pairwise interaction models like Lotka-Volterra provide an important and flexible foundation, but notably absent is an explicit mechanism mediating interactions. Consumer-resource models incorporate mechanism, but describing competitive and mutualistic interactions is more ambiguous. Here, we bridge this gap by modeling a coarse-grained version of a species’ true, cellular metabolism to describe resource consumption via uptake and conversion into biomass, energy, and byproducts. This approach does not require detailed chemical reaction information, but it provides a more explicit description of underlying mechanisms than pairwise interaction or consumer-resource models. Using a model system, we find that when metabolic reactions require two distinct resources we recover Liebig’s Law and multiplicative co-limitation in particular limits of the intracellular reaction rates. In between these limits, we derive a more general phenomenological form for consumer growth rate, and we find corresponding rates of secondary metabolite production, allowing us to model competitive and non-competitive interactions (e.g., facilitation). Using the more general form, we show how secondary metabolite production can support coexistence even when two species compete for a shared resource, and we show how differences in metabolic rates change species’ abundances in equilibrium. Building on these findings, we make the case for incorporating coarse-grained metabolism to update the phenomenology we use to model species interactions.

## Introduction

A central goal in ecology is to understand and predict the dynamics in communities of interacting species (Holt, 1977; Loreau, 2010; Vellend, 2010, 2016). Mathematical models allow us to generate and test theoretical predictions, and the development of such models leads to a hierarchy of challenges. The first challenge is determining an appropriate functional form describing species dynamics. A range of functional forms with increasing complexity has been used and each has strengths and weaknesses which are often context dependent (Holling, 1959; Abrams, 1982; DeAngelis et al., 1989; Murdoch et al., 2003; Mougi and Kondoh, 2012). Second, we need to parameterize these equations, for example by quantifying the strength and sign of species interactions in a given environmental context. While some attempts have been made to parameterize natural systems, accurately fitting interaction strengths remains challenging in both empirical and theoretical work (Schoener, 1983; Tilman, 1987; Ives et al., 2003; Marino et al., 2014; Carrara et al., 2015; Terry et al., 2017; Barner et al., 2018). Finally, we may wish to determine how changes in the environmental context will modify species interactions, dynamics, and even coexistence. Integrating each of these goals will lead to the development of robust models which can predict the dynamics of complex communities, even when the environmental landscape changes within and across ecosystems.

There are two primary approaches to mathematically model the dynamics of interacting species. The Lotka-Volterra equations characterize species interactions in terms of the net, direct effect of one population on another’s growth rate (Lotka, 1932; Volterra, 1926, see Supplemental). However, because these models lack an explicit description of the mechanisms mediating interactions, it is often difficult to translate the inferred interactions in different environmental contexts (Abrams, 1983; Grilli et al., 2017). For example, if two species compete, it is often because they consume common resources (Gause and Witt, 1935; MacArthur, 1970; Schoener, 1983). But, these models assume that resource dynamics can be safely ignored because resource dynamics are faster than consumer dynamics (MacArthur, 1970). This exposes an important context-dependence of Lotka-Volterra type equations: the strength and even the sign of a pairwise interaction may depend on what resources are present (Xiao et al., 2017). An alternate approach, consumer-resource equations, characterizes competitive interactions as the explicit result of shared resource consumption (Tilman et al., 1982; Grover, 1990; Litchman, 2003; Abrams, 2009, see Supplemental). These models produce species interactions as an emergent property dependent on shared resource consumption. Assuming we can infer or otherwise estimate consumer preferences, one critical aspect of the environmental context is now explicitly characterized, via resource input rates. As such, species dynamics across resource landscapes can be understood better than in the case of Lotka-Volterra, where the effect of the environment is implicit (Tilman, 1977; Grover, 1990, 2011).

The more explicit mechanism in a consumer-resource model gives an advantage, but there is a cost. Lotka-Volterra equations are extremely flexible and can straightforwardly incorporate a mixture of antagonistic and mutualistic interactions simply by altering the signs of entries in the community matrix (Mougi and Kondoh, 2012; Allesina and Tang, 2012). But for consumer-resource models we have to be careful about how those mechanisms are formulated. Consumption can take a variety of forms depending, for example, on whether resources are substitutable or essential (Tilman, 1980), and mutualistic interactions can occur via resource production and exchange (i.e., cross-feeding) (Loreau, 2001; Freilich et al., 2011; Zelezniak et al., 2015; Sun et al., 2019). The latter in particular is underexplored relative to consumption (Butler and O’Dwyer, 2018), and an open question is to what extent the precise form of exchange might affect community dynamics and species coexistence.

So how do we retain the advantages of consumer-resource models, but also incorporate a mixture of species interactions? Current consumer-resource models are largely agnostic to what happens inside cells or organisms. For many systems, this approach may be valid especially when the resources (i.e., prey) belong to higher trophic levels, self-regulate, and/or closely match the stoichiometric requirements of the consumer (Sterner and Elser, 2002; Cherif and Loreau, 2007; Hall, 2009). However, this may not be valid for microorganisms. First, most microorganisms consume abiotic resources which do not self-regulate. Second, any single resource generally does not satisfy the full stoichiometric requirements of metabolism. For example, heterotrophic microorganisms require organic carbon, but they still require nitrogen, phosphorus, and other resources to grow and reproduce. Because growth depends on multiple resources, population dynamics may depend on a single limiting resource (e.g., Liebig’s Law of the Minimum: von Liebig and Gregory, 1842; Odum, 1959) or an interaction between resources (e.g., multiplicative co-limitation: Harpole et al., 2011). Likewise, the consumption and transformation of resources depends on how cells produce biomass, energy, and metabolic byproducts, and species interactions may therefore depend on metabolic rates and the release of metabolic byproducts.

Here, we propose that going one level deeper into cellular metabolism will allow us to generalize consumer-resource models in a meaningful way and give them the same flexibility to describe multiple interaction types as models of direct species interactions. First, we introduce a general metabolically informed consumer-resource model for a single species. In this model, we include internal cellular dynamics via a simplified metabolic network, which require less knowledge of a particular species’ idiosyncrasies but still captures the major metabolic events transforming resources. Using this model, we explore population dynamics in variable resource environments. Next, we expand the approach to incorporate a second species. Here, both species share a common resource, but the second species also uses a metabolic byproduct produced by the first species. As such, our model includes competition for the shared resource and facilitation via metabolic cross-feeding. We then explore how the resource landscape and internal metabolic rates interact to demonstrate how competition and facilitation mediate species dynamics and coexistence conditions under different resource use scenarios. In summary, our findings demonstrate how a metabolically informed consumer-resource model for a single species can produce both Liebig’s Law and multiplicative co-limitation in particular limits of the metabolic rates. Furthermore, when a second species is included we expand the expectations of Liebig’s Law and multiplicative colimitation to include cross-feeding and demonstrate both species dynamics and equilibrium abundances across resource landscapes.

## Methods and Results

### The Metabolically Informed Consumer-Resource Model

We know a substantial amount about the internal physiology of microorganisms, and there have been large advancements in the development of flux balance models which use biochemistry and genomics to describe (to some level of approximation) every reaction that occurs within a cell (Kauffman et al., 2003; Orth et al., 2010). Likewise, models based on the dynamic energy budget theory provide a complete mass and energy balance approach for molecular processes, cellular physiology, and population growth (Kooijman, 2001; Nisbet et al., 2008). More recently, flux balance models have been applied in the context of entire microbial communities and their interactions (Embree et al., 2015; Zomorrodi and Segrè, 2016; Pacheco et al., 2019). However, we propose that including a full description of a metabolic network may not be required to develop a useful ecological model. Simplified metabolic models have been used to understand metabolic partitioning (Kempes et al., 2012) and trophic tradeoffs (Chakraborty et al., 2017). Furthermore, models that add some intracellular dynamics, such as Droop’s model which accounts for cell quotas and intracellular resource storage (Droop, 1974), have been used to better understand how consumers contend with nutrient limitations and how trait trade-offs underlie coexistence (Cherif and Loreau, 2010; Litchman et al., 2015). Here, we model the internal dynamics of a microorganism via simplified metabolic networks, which require less knowledge of the particular species’ idiosyncrasies but still capture the major metabolic events transforming resources. Our prototypical metabolic model is based on a basic fermentation reaction, homolactic fermentation, which uses glucose and phosphate and produces lactate (Figs. 1 & S1). In this reaction, glucose and phosphate are used to form new biomass, and glucose is used to produce chemical energy via fermentation. The energetic component results in the production of a byproduct, lactate, which is exported back into the environment (i.e., excretion); therefore, efficiency is emergent property determined by the balance between the biomass and energy production.

**Fig. 1.**
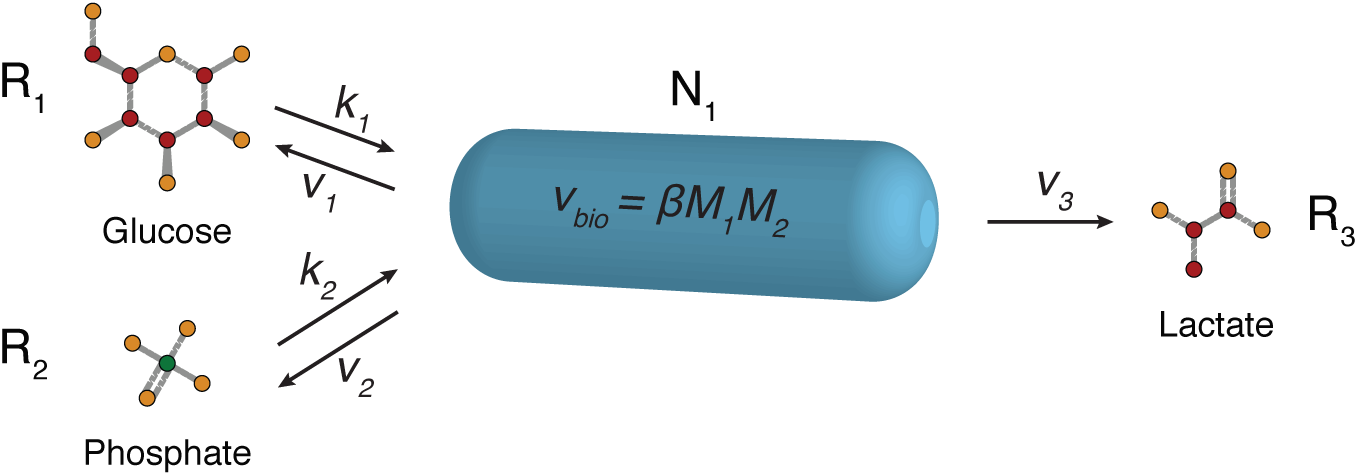
Conceptual Model. Our metabolically informed consumer-resource model uses fermentation as a prototype. Fermentation is an anaerobic—usually sugar consuming—metabolic lifestyle, and is the primary anaerobic energy-producing reaction for many microorganisms (Gottschalk, 1986). A signature of fermentation is that it results in byproducts such as organic acids, alcohols, and/or gases, which are produced due to the incomplete resource oxidation during the energy producing reactions. When organic acids (e.g., lactate) are produced, they are often used as resources by other microorganisms. In our model, we simplify the true biochemistry (see Supplemental, Fig. S1) by assuming glucose (*R*_1_) and phosphate (*R*_2_) enter the cell at uptake rates *k*_*i*_, biomass (*N*_1_) is generated at rate *βM*_1_*M*_2_, and the metabolic byproduct, lactate, is exported from the cell at rate *ν*_3_. In addition, we assume that any unused glucose and phosphate are exported at rates *ν*_1_ and *ν*_2_.

Here, we assume that cellular metabolism relies on the interaction of sugar and phosphate, producing new biomass and a byproduct (i.e., lactate).

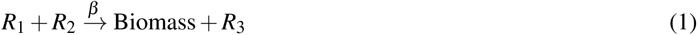

*R*_1_ here is the sugar, *R*_2_ represents a source of phosphorus, and *R*_3_ is the metabolic byproduct lactate. We model this intracellular reaction as occurring at rate *β*. Next, we assume that the organism grows in a chemostat-type environment, where resource inflow rates are held constant. In addition, we assume that metabolic rates are not limited by any other factors and that resources are not toxic at high concentrations and therefore do not inhibit growth. Given that we consider one phosphate molecule (i.e., we assume 1:1 stoichiometry), this already simplifies the resource requirements relative to the true process. But we will see general principles emerge.

We now assume that intracellular processes are always close to equilibrium, so that we can balance fluxes for the internal cellular densities of the two input resources (labeled *M*_1_ and *M*_2_). This leads to:

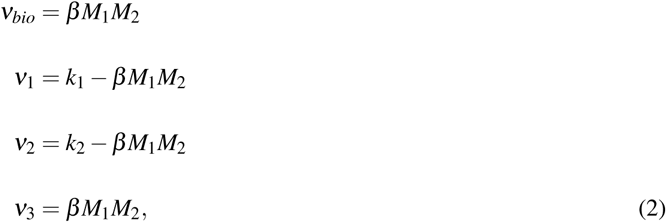

for uptake rates *k*_1_ and *k*_2_, which can depend in an arbitrary way on external resource concentrations *R*_1_, *R*_2_, and *R*_3_, outflow (i.e., export) rates *ν*_1_, *ν*_2_, and *ν*_3_, and biomass production *ν*_*bio*_ (Fig. 1). We make a natural assumption that export of each resource is determined by passive excretion: i.e. that *ν*_*α*_ = *λ*_*α*_ *M*_*α*_ for each metabolite, where *λ*_*α*_ is a species- and resource-specific constant. This resource excretion, or leakiness, prevents unbounded internal resource accumulation, but it can also mediate species interactions (e.g., cross-feeding) and therefore structure communities (Pfeiffer and Bonhoeffer, 2004; Morris, 2015; Sun et al., 2019). As such, the outflow rates (*ν*_*α*_) represent *use it or lose it* internal resource dynamics and therefore intracellular resources that are not used are exported back into the external environment.

We can solve this system of polynomial equations (Eq. 2) for internal resource concentrations to obtain:

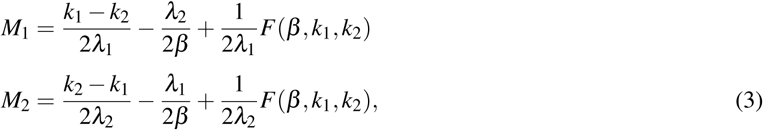

while biomass production is given by:

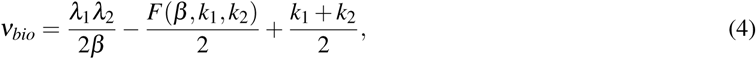

and where 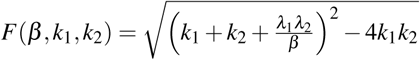 depends on uptake, export, and reaction rates.

Using these intracellular concentrations, we can now develop a set of generalized consumer-resource equations using uptake rates and this simplified flux balance analysis (Eq. 4) as the only building blocks. First, we focus on one species and model its uptake rate *k*_*α*_ of resource *α* as *C*_*α*1_*R*_*α*_, where *R*_*α*_ is the *external* (environmental) concentration of this resource. Applying this assumption, we derive a general set of equations for the consumer and three resources:

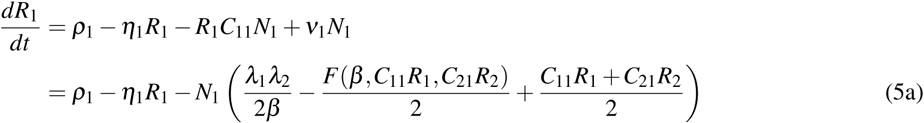

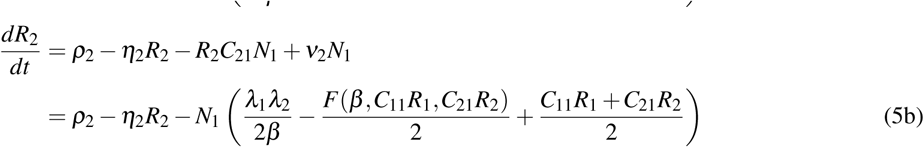

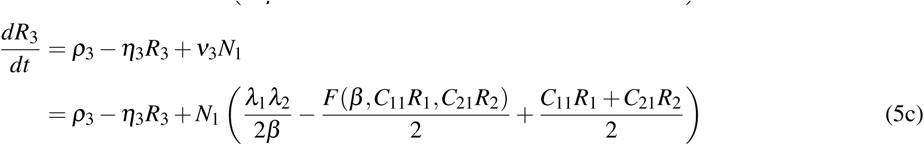

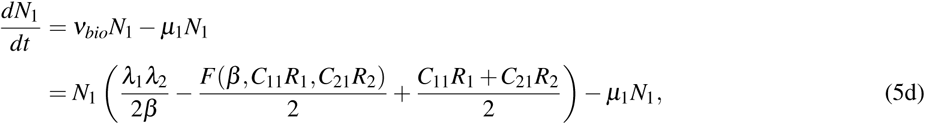

where *ρ*_*α*_ and *η*_*α*_ are inflow and outflow rates for each of the three resources, and *µ*_1_ is the mortality rate of the consumer (see Table 1). For the resources (Eqs. 5a–5c), the extracellular resource concentrations are determined by the inflow and outflow rates minus the consumer-density dependent components which include uptake, leakage, and conversion into biomass (Eq. 4). For the consumer (Eq. 5d), the production of new biomass is determined by the density dependent uptake, leakage, and conversion of resources into biomass (Eq. 4) minus the density dependent mortality. Here, we have assumed that uptake, *k*_*α*_, increases linearly with resource concentration, but this could easily be modified to saturate using Monod dynamics by changing *k*_*α*_ to (*C*_*α*1_*R*_*α*_)*/*(*K*_*α*1_ + *R*_*α*_), where *K*, the half-saturation constant, is a species- and resource-specific constant.

**Table 1.** Model Notation. Note: resource concentrations are given in cell equivalents – moles required to generated one cell.

| Symbol | Meaning | Units |
| --- | --- | --- |
| $N_i$ | Number of cells in population $i$ | cell |
| $R_\alpha$ | Concentration of resource $\alpha$ | cell equivalent |
| $\rho_\alpha$ | Inflow rate of resource $\alpha$ | cell equivalent $\text{hr}^{-1}$ |
| $\eta_\alpha$ | Outflow rate of resource $\alpha$ | cell equivalent $\text{hr}^{-1}$ |
| $C_{\alpha i}$ | Uptake rate of resource $\alpha$ by population $i$ | cell equivalent $\text{hr}^{-1}$ cell$^{-1}$ |
| $v_i$ | Mortality rate of population $i$ | cell $\text{hr}^{-1}$ |
| $\lambda_\alpha$ | Leakage rate of resource $\alpha$ | cell equivalent $\text{hr}^{-1}$ |
| $\beta_i$ | Internal metabolic rate of population $i$ | cell $\text{hr}^{-1}$ |

We now note two limits of the simplified flux balance analysis (Eqs 3–4). The first limit is where *β* (i.e., the internal metabolic rate) is large relative to the other rates,

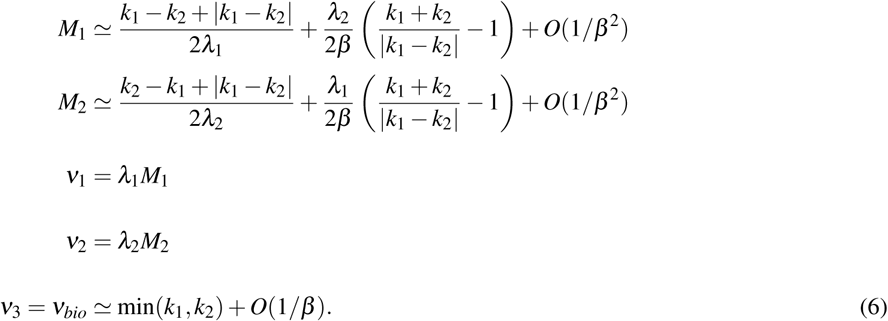

Where *O*(1*/β*) represents the linear approximation of the higher order terms of the Taylor Series expansion. Note that we have to keep the *O*(1*/β*) terms for *M*_1_ and *M*_2_, because for at least one of the two (depending on whether *k*_1_ > *k*_2_ or *k*_1_ < *k*_2_) the *O*(1*/β*) term vanishes in this limit of fast reaction rate *β*. Therefore, when *β* is large relative to the other rates the consumer-resource equations become:

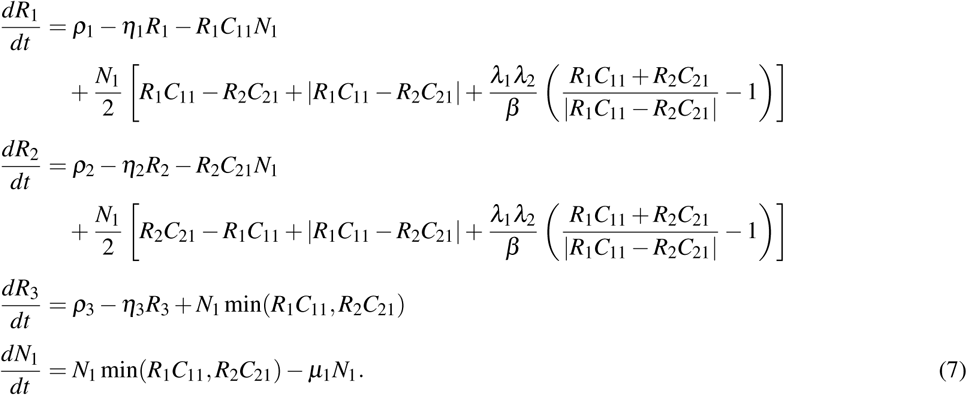

Hence, we recover Liebig’s law (i.e. the net growth rate of the consumer is min(*R*_1_*C*_11_, *R*_2_*C*_21_)). Using these equations in numerical simulations, we observe that the consumer abundance saturates (Fig. 2A), and the final abundance depends on the inflow rate of the more limiting resource (Fig. 2B). As such, the model in this limit behaves similar to classical consumer-resource models. In addition, we generate novel terms for the byproduct production rate, which in this fast reaction rate limit is ≃ *N*_1_ min(*R*_1_*C*_11_, *R*_2_*C*_21_), and we find corresponding equations for the uptake and export of glucose and phosphate.

**Fig. 2.**
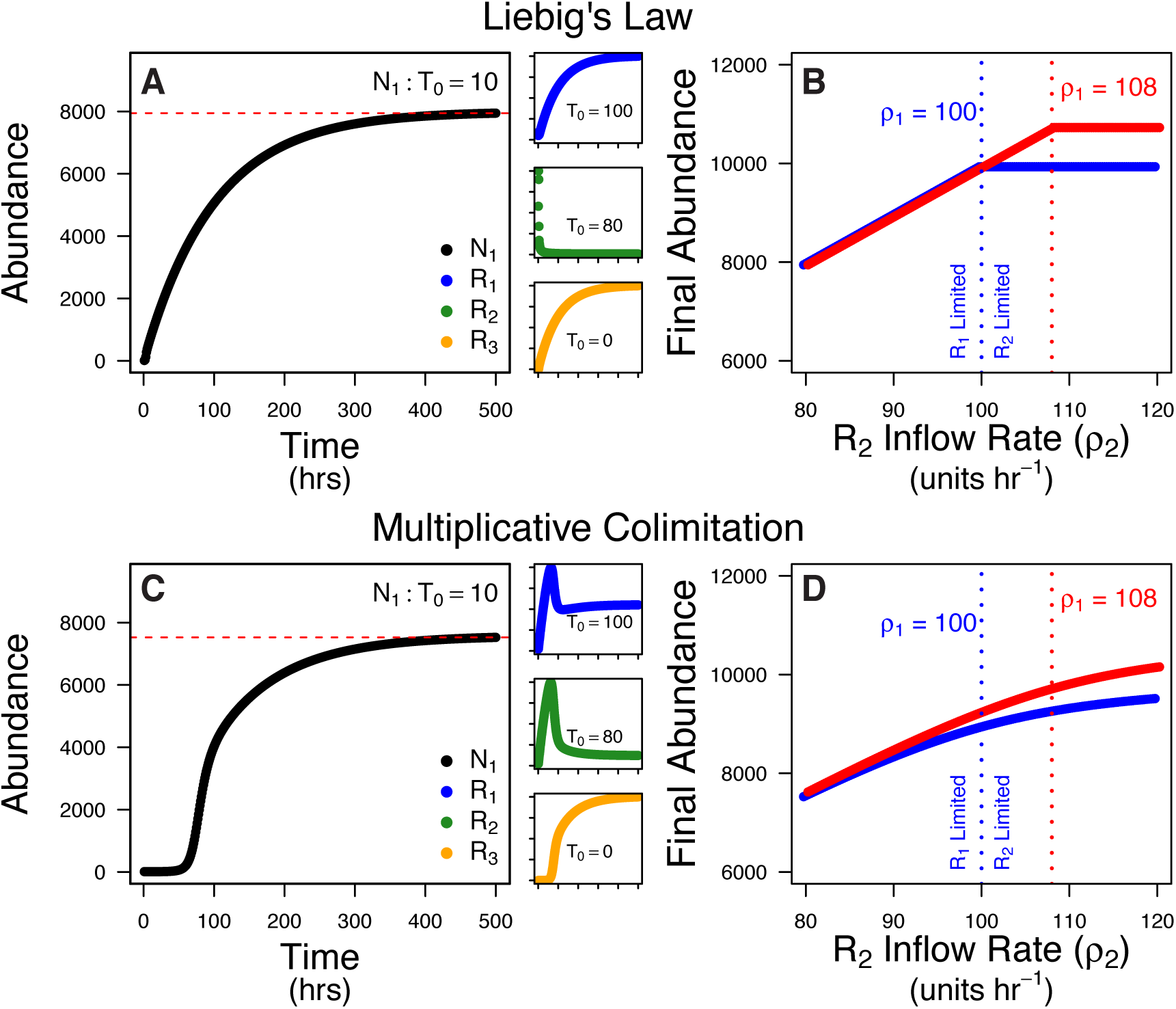
Single Species Simulation. We used numerical simulations of our single-species model to determine how the resource landscape changes population dynamics. In our model, *N*_1_ is the population, *R*_1_ and *R*_2_ are the consumed resources (i.e., sugar and phosphate), *R*_3_ is a metabolic byproduct, *ρ* is the resource inflow rate, and *β* is the internal metabolic rate of the consumer. **A & B**: In the limit where *β* is high relative to other rates (Eq. 7), we recover Liebig’s Law of the Minimum. Consumer abundance saturates, but the final abundance depends on the inflow rate, *ρ*_*α*_, of the more limiting resource. When the inflow of *R*_2_ (*ρ*_2_) is lower than the inflow of *R*_1_ (*ρ*_1_), *R*_2_ becomes depleted. At saturation, *R*_1_ is in excess while *R*_2_ is limiting (**A**). Changes in the inflow rates alters the final abundance (**B**), but the final abundance will always be determined by the more limiting resource. When *ρ*_1_ = 100 (blue) the final abundance will increase until *ρ*_2_ = 100 (blue dashed line), and when *ρ*_1_ = 108 (red) the final abundance will increase until *ρ*_2_ = 108 (red dashed line). In contrast, in the limit where *β* is low relative to other rates (Eq. 9), we recover multiplicative co-limitation. Consumer abundance still saturates, albeit at a lower final abundance given the same resource inflow rates and with a growth lag-phase (**C**). As consumer abundance increases, both *R*_1_ and *R*_2_ are depleted. In addition, the final abundance increases as inflow rates increase (**D**). When *ρ*_1_ = 100 (blue) the final abundance will continue to increase past *ρ*_2_ = 100, and when *ρ*_1_ = 108 (red) the final abundance will continue to increase past *ρ*_2_ = 108. Simulations based on Eq. 5 using the following parameters: *η*_*α*_ = 0.01, *C*_*α*1_ = 0.01, *µ*_1_ = 0.01, *λ*_*α*_ = 0.1, *β*_1_ = 10 (**A**) or 10^−6^ (**B**), *ρ*_1_ = 100 or 108, and *ρ*_2_ varies between 80 and 120 (**A** & **C**: *ρ*_1_ = 100, *ρ*_2_ = 80). Resource concentrations are expressed in cell equivalents (i.e., moles required to generate a cell) and we assume 1:1 resource stoichiometry.

The second limit is where *β* (i.e., the internal metabolic rate) is small relative to the other rates,

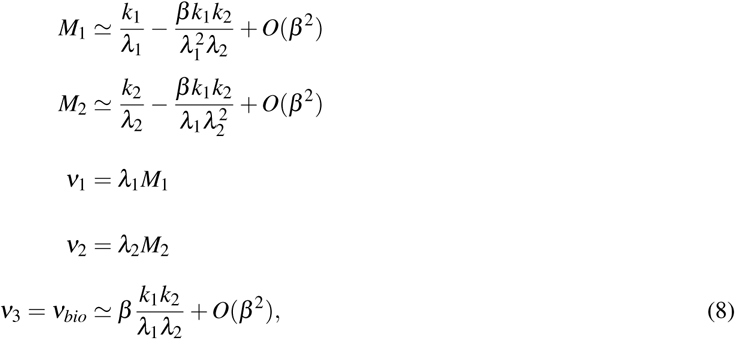

so that the consumer-resource equations become:

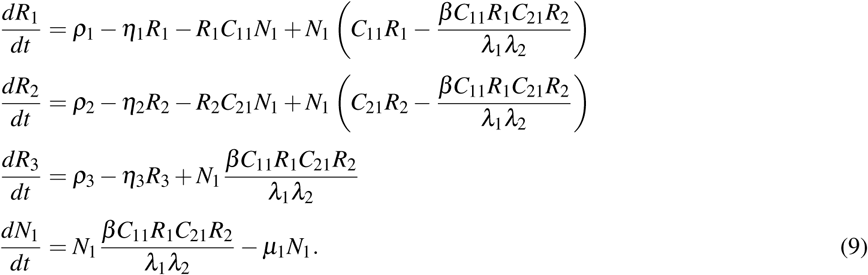

Hence, we recover multiplicative co-limitation by the two resources (i.e., the net growth rate of the consumer is 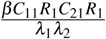). Using numerical simulations, we observe that consumer abundance, *N*_1_ saturates as expected but now includes a growth lag-phase (Fig. 2C). The lag-phase (i.e., period with no net population growth: 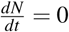) occurs while 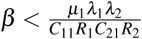. When *R*_1_ and *R*_2_ accumulate (Fig. 2C), the denominator increases and the inequality flips which yields population growth. Due to the parameters used in the simulation above, the equation for the lag-phase simplifies to 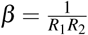; therefore, our simulation demonstrates how the lag-phase is controlled by an interaction between the resource environment and the internal metabolic rates. Finally, the final abundance depends on the inflow rate of both resources (Fig. 2D). An increase in the inflow rate of either resource will yield a higher final population abundance.

In summary, from this coarse-grained representation, we recover two classic outcomes of consumer-resource theory by taking limits of the internal reaction rate *β*. We can also generalize these classic limits for intermediate *β*, in a way that is not currently used in consumer-resource models and falls neither into the category of Liebig’s Law nor multiplicative co-limitation. Finally, we find functional forms for the production rate of metabolic byproducts (and excretion of other resources) in each of these limits and in between. Our model therefore demonstrates how we can generalize the functional form of consumer-resource models by considering realistic, simplified intracellular processes.

### The Two Species Model

We now modify the model above to incorporate a second species. Here, both species use *R*_2_ (phosphate), but the second species, *N*_2_, uses a combination of *R*_2_ and *R*_3_ (lactate) to generate new biomass (Fig. 3). While we are using this as a model with both competition (e.g., shared resources) and facilitation (e.g., metabolic cross-feeding) interactions, it also represents the natural cross-feeding interaction between organisms. For example, the model could represent the interactions between the lactate producing (e.g., *Bifidobacterium adolescentis*) and lactate consuming (e.g., *Eubacterium hallii*) bacteria found in human and animal digestive systems (Duncan et al., 2004). Likewise, other examples of these interactions would include the exchange of vitamin precursors and amino acids by auxotrophic organisms (Carini et al., 2014; Embree et al., 2015). These metabolic cross-feeding interactions are common in microbial systems (Pfeiffer and Bonhoeffer, 2004; Morris et al., 2012; Mee et al., 2014; Morris, 2015; Tasoff et al., 2015; Zelezniak et al., 2015; Sun et al., 2019) and have industrial applications (Jiao et al., 2012). Cross-feeding has been shown to promote species diversity and increase net ecosystem production (Morris et al., 2012; Pande et al., 2014). In our model, productivity (and efficiency) increase due to the second species using the metabolic byproducts of the first (Fig. 3). As such, our model demonstrates how competition (i.e., shared resources) and facilitation (i.e., production and consumption of intermediate resources) mediate species dynamics and coexistence conditions and can be used to understand natural and engineered microbial systems.

**Fig. 3.**
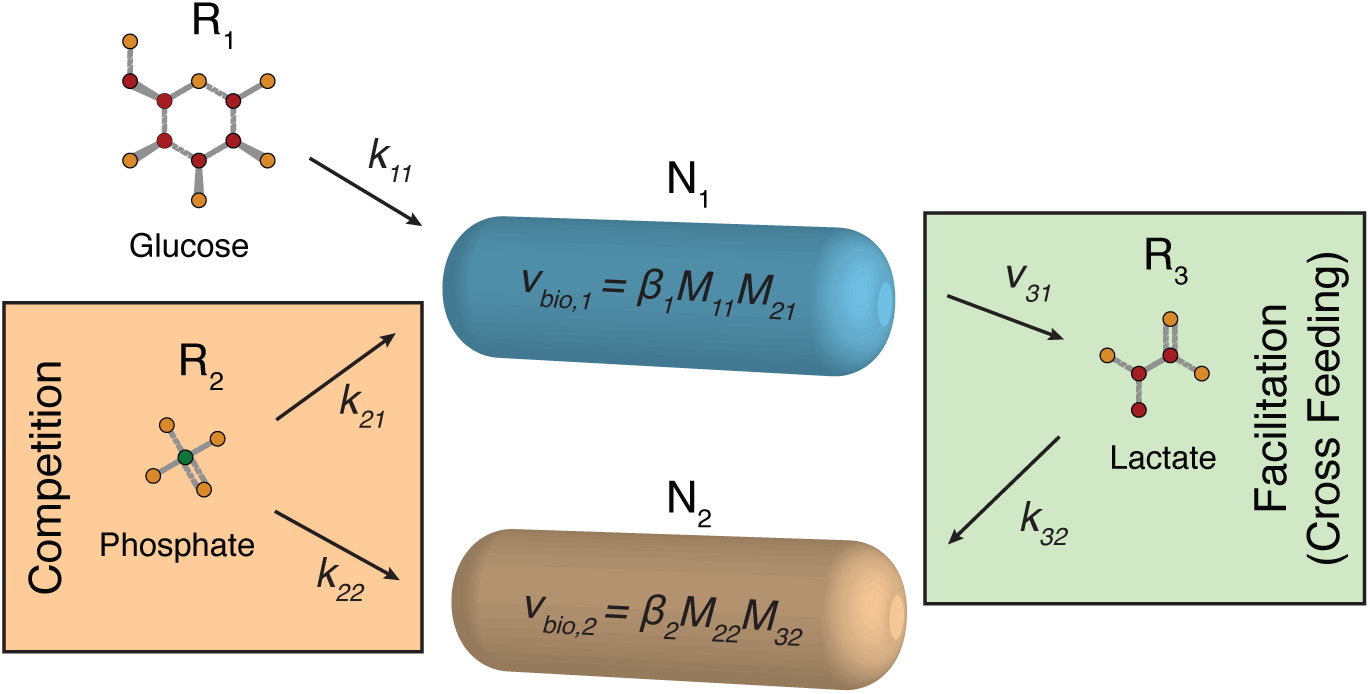
Two Species Model. In the two species model, two species interact through competition and facilitation. For each species, we use a simplified metabolic networks to describe resource uptake rates (*k*_*αi*_), biomass generation rates (*β*_*i*_*M*_*αi*_*M*_*αi*_), and metabolic byproduct export rates (*ν*_*αi*_) for species *i* and resource *α*. Species *N*_1_ consumes glucose (*R*_1_) and phosphate (*R*_2_) and produces lactate (*R*_3_). Species *N*_2_ competes for phosphate (*R*_2_) but also uses the lactate produced by species *N*_1_. In addition, all internal resources (byproducts and unused resources) are exported at rate *ν*_*α*_ (not shown). As such, facilitation is modeled by cross-feeding (i.e., production and consumption of lactate) between species *N*_1_ and *N*_2_.

First, we define two distinct internal metabolic processes, one for each consumer species:

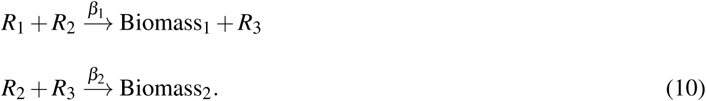

Consumer *N*_2_ may also produce a metabolic byproduct, but we have not included such a process here because we are focusing on competition for *R*_2_ and facilitation through the production of *R*_3_ by species *N*_1_ (Fig. 3). Importantly, we now have two internal reaction rates, *β*_1_ and *β*_2_. Here we focus on how these rates, both relative to each other and also to the other rates in the model, affect species coexistence. This approach allows us to use the coarse-grained metabolic processes as the mechanisms underlying species interactions. Furthermore, it allows us to explore the potential for changes in species coexistence due to metabolic (i.e., reaction rates) and landscape (i.e., inflow rates) factors.

Generalizing the approach in the previous section, and again assuming internal flux balance, we can define the fluxes for an individual belonging to species *N*_1_ as:

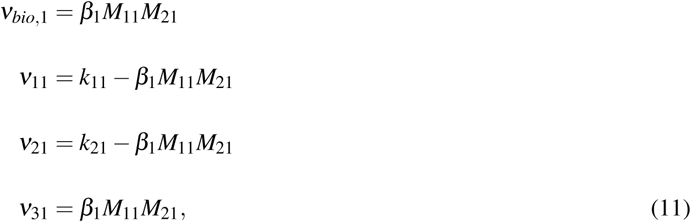

for internal concentrations *M*_*α*1_ and uptake rates *k*_*α*1_. While for an individual of species *N*_2_ we have:

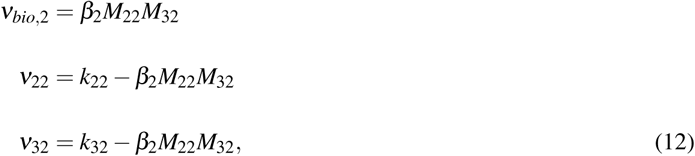

for internal concentrations *M*_*α*2_ and uptake rates *k*_*α*2_. We also assume that all resources can be excreted from both consumers, but to simplify the model slightly we will assume that the specific export rates are equal, *λ*, broadly consistent with passive diffusion across sufficiently similar cell wall types. Based on these assumptions, we can solve for internal equilibrium in both cell types to obtain:

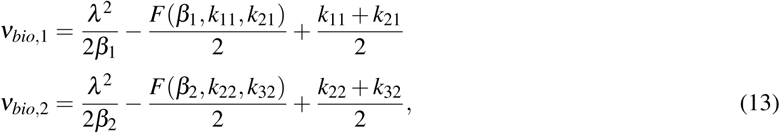

where, similarly to the one species case, the function 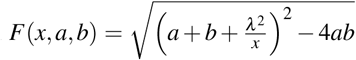 (see: Eqns: 3 & 4). Finally, we can put all of this together to generate a set of equations for the dynamics of both populations and the external concentrations of all three resources.

To focus on the effects of internal reaction rates and the resource landscape, we will further simplify our model by making a few assumptions. We will assume that the outflow rates for each resource are the same, so that *η*_*i*_ = *η*, and that the per capita mortality rates for each consumer are equal, so that *µ*_*i*_ = *µ*. We will also again assume that the per capita uptake rate of resource *α* by species *i* can be written as *C*_*αi*_*R*_*α*_. We will then determine the effects of internal reaction rate by independently changing *β* (i.e., the internal metabolic rate) for each species. In addition, we will change the landscape conditions by exploring the inflow rates for each resource *ρ*_*α*_. Given these assumptions, our general two species consumer-resource model is:

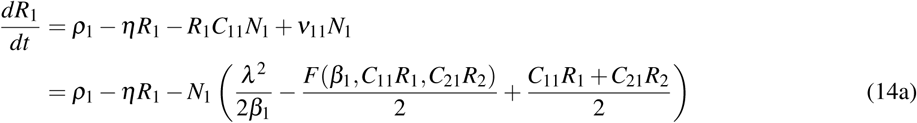

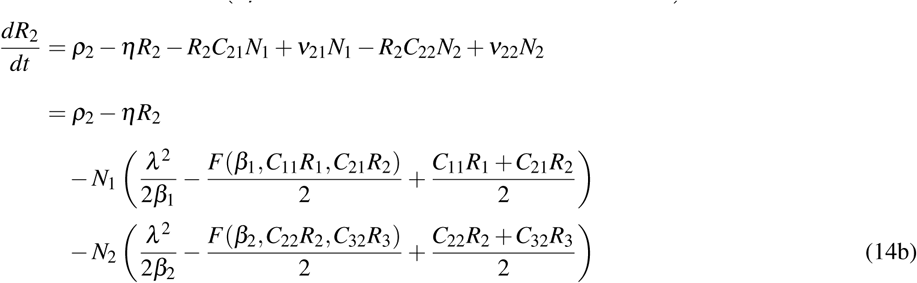

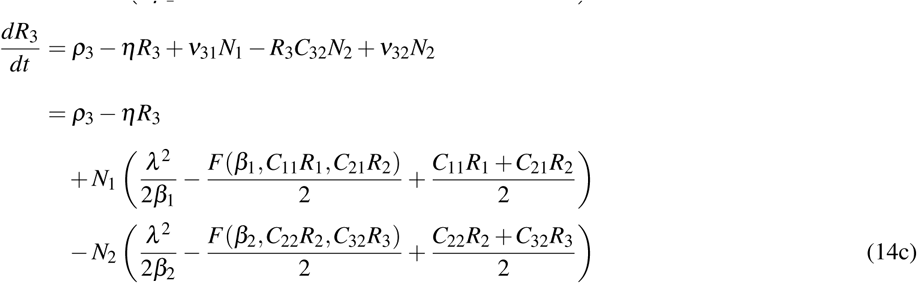

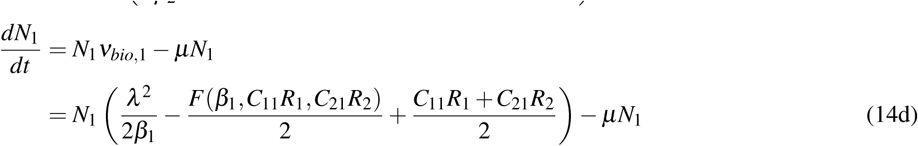

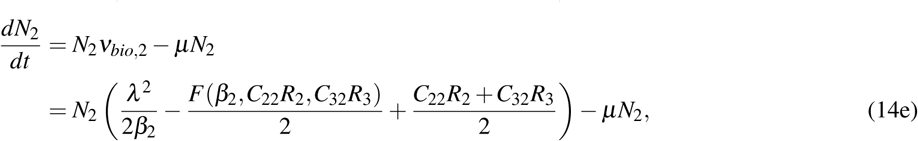

where *ρ*_*α*_ and *η*_*α*_ are inflow and outflow rates for each of the three resources, and *µ*_1_ is the mortality rate of the consumer (see Table 1). For the resources (Eqs. 14a–14c), the extracellular resource concentrations are determined by the inflow and outflow rates minus the consumer-density dependent components of each consumer which include uptake, leakage, and conversion into biomass (Eq. 13). For the consumers (Eqs. 14d & 14e), the production of new biomass is determined by the density dependent uptake, leakage, and conversion of resources into biomass (Eq. 13) minus the density dependent mortality.

Using numerical simulations, we model the resource and consumer dynamics and determine equilibrium conditions (Figs. 4 & 5). First we consider when the internal metabolic rates, *β*_*i*_, are the same. When internal metabolic rates are both high, species coexist at a density determined by the shared resource inflow rate (i.e., *ρ*_2_) until the inflow rate of the unshared resource, *R*_1_, exceeds the inflow rate of the shared resource, *R*_2_. When *ρ*_1_ is greater than *ρ*_2_ (i.e., *ρ*_1_*/ρ*_2_ > 1), species *N*_1_ will outcompete species *N*_2_ for *R*_2_, and *N*_2_ will become rare (Fig. 4A). In our simulations, species *N*_2_ remains in the system with rare abundance due to resource leakage from species *N*_1_, but would likely go extinct due to stochastic fluctuations, and so coexistence in this limit is somewhat tenuous. These findings expand the expectations of Liebig’s Law to two cross-feeding species and demonstrate both species dynamics and equilibrium abundances across various resource landscapes. In contrast, when internal metabolic rates are low, species relative abundances are determined by their respective required resources based on multiplicative co-limitation and both species now demonstrate growth lag-phases. These findings expand the expectations of multiplicative co-limitation to two cross-feeding species. However, since species *N*_1_ will not be resource limited when *ρ*_1_ is greater than *ρ*_2_, then species *N*_2_ will maintain a higher relative abundance across wider resource landscape (Fig. 4B, see Eq. 9). Together, we find that if internal rates are the same but either high or low compared to the other rates in our model, then our results expand the expectations of Liebig’s Law and multiplicative co-limitation to the two-species system with a metabolic dependency. Between these limits, we find a smooth transition between the two scenarios as coexistence conditions become decoupled from classical resource limitation predictions when the internal metabolic rates decrease (Fig. 4C). In addition, we find that coexistence depends on both the internal metabolic rates and the resource inflow rates even when uptake rates are the same. Furthermore, while species *N*_2_ depends on species *N*_1_ for the production of *R*_3_, external inputs of *R*_3_ (i.e., *ρ*_3_ > 0) alone are not enough to allow species *N*_2_ to outcompete species *N*_1_ but will result in a lower final density (see Supplemental).

**Fig. 4.**
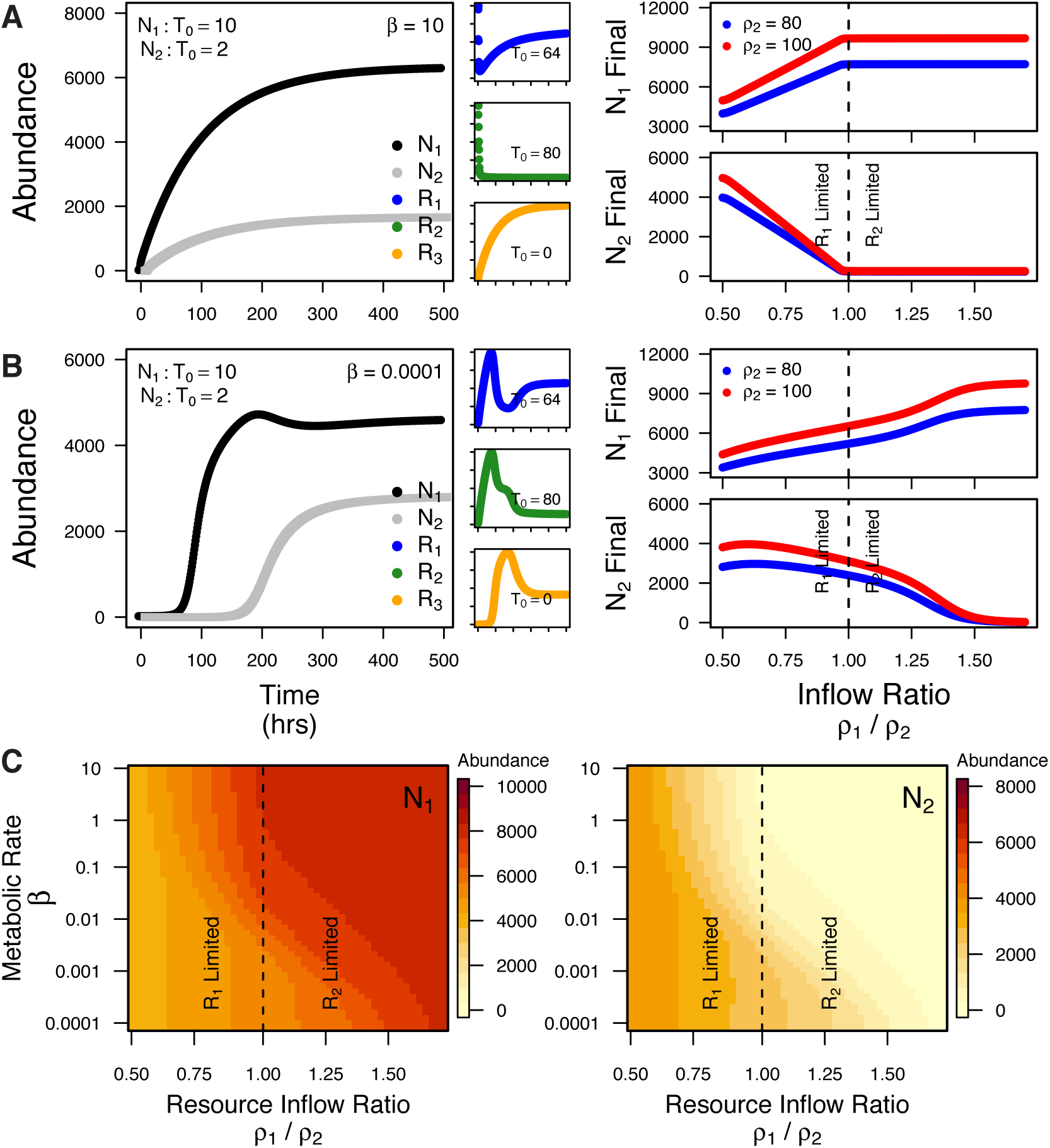
Two Species with the Same Metabolic Rates. We simulated species dynamics in the two-species model with equal internal metabolic rates, *β*_*i*_. In this model, two species compete for a shared resource (*R*_2_), but *N*_1_ consumes *R*_1_ and produces *R*_3_ as a metabolic byproduct which is consumed by *N*_2_. When metabolic rates are high, species abundances saturate and species can coexist provided the appropriate resource conditions (**A**). When the shared resource inflow rate (*ρ*_2_) is higher than the inflow rate of *R*_1_ (*ρ*_1_), then the two species coexist at a relatively high abundance because *N*_1_ is more limited by its exclusive resource (*R*_1_). However, when *ρ*_1_ is higher than *ρ*_2_ (i.e., *ρ*_1_*/ρ*_2_ > 1), then *N*_1_ can outcompete *N*_2_ for the shared resource (*R*_2_) and the final abundance of *N*_2_ will be reduced. Even when the shared resource inflow rate is increased (*ρ*_2_ = 100 vs. 80), the final abundance of *N*_2_ does not increase when *ρ*_1_*/ρ*_2_ > 1. When metabolic rates are low, species still coexist, but the range of coexistence is greatly expanded (**B**). While the final abundance of *N*_2_ still decreases as the resource inflow ratio (*ρ*_1_*/ρ*_2_) approaches 1, it does not decrease at the same rate and the final abundance remains relatively high even when the resource inflow ratio is > 1. In this simulation, the resource inflow ratio needs to be greater than around ∼ 1.5 before *N*_1_ outcompetes *N*_2_ and *N*_2_ becomes rare. Between these metabolic rates, there is a smooth transition (**C**). When both resources have equal inflow rates, the final abundance of *N*_1_ decreases and *N*_2_ increases as *β* drops below 1. Simulations based on Eqs. 14a–14e using the following parameters: *η*_*α*_ = 0.01, *C*_*αi*_ = 0.01, *µ*_*i*_ = 0.01, *λ*_*α*_ = 0.1, *β*_*i*_ = 10 (high) or 10^−6^ (low), *ρ*_2_ = 80 or 100, and *ρ*_1_*/ρ*_2_ varies between 0.5 and 1.7 (**A**_*left*_ & **B**_*left*_:*ρ*_1_ = 64, *ρ*_2_ = 80). Resource concentrations are expressed in cell equivalents (i.e., moles required to generate a cell) and we assume 1:1 resource stoichiometry.

**Fig. 5.**
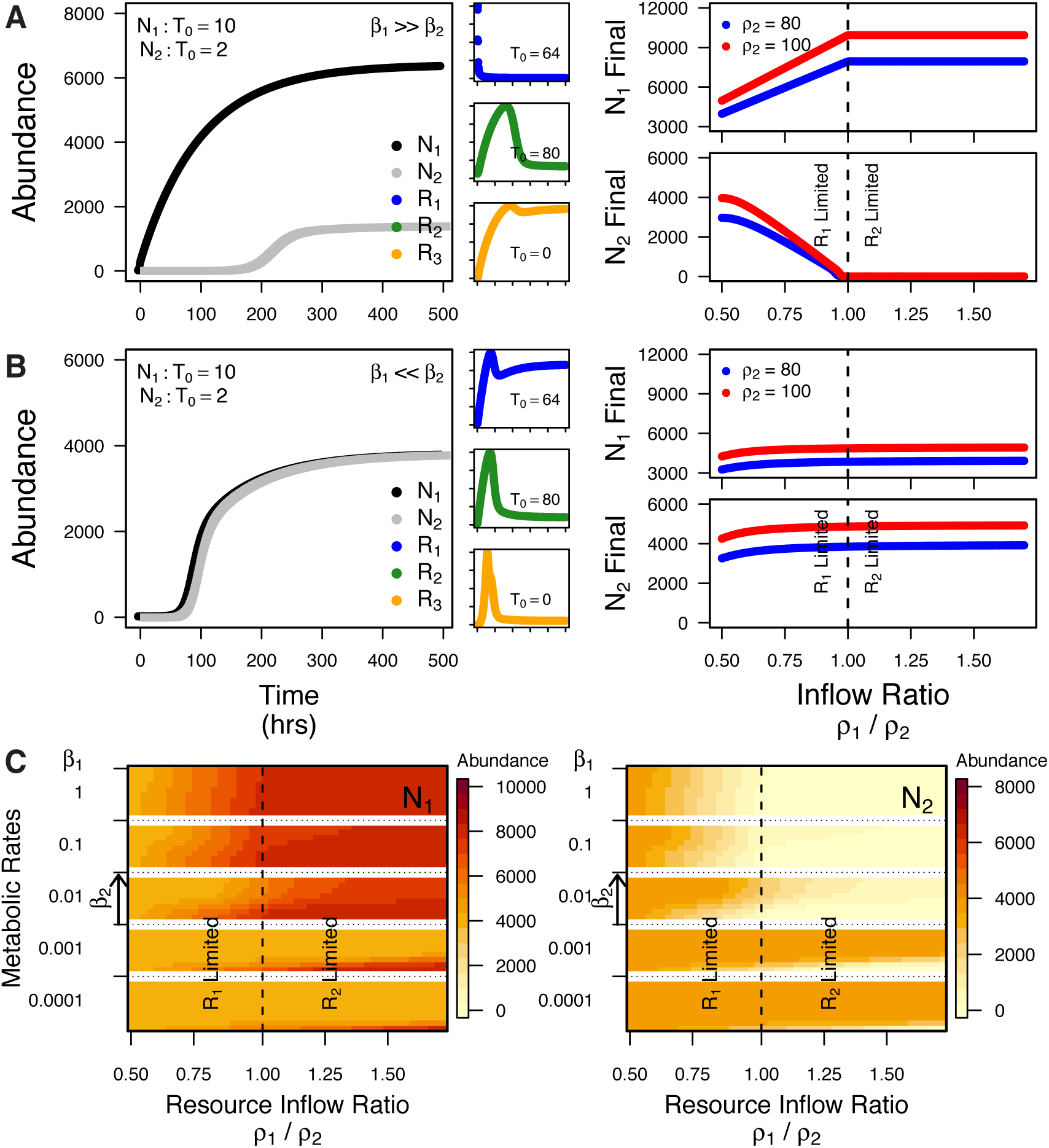
Two Species with Different Metabolic Rates. We simulated species dynamics in the two-species model where the species have different internal metabolic rates, *β*_*i*_. In this model, one of the species has high and the other low metabolic rates compared to the other rates in the model. The species compete for *R*_2_, and species *N*_1_ consumes *R*_1_ and produces *R*_3_ which is used by species *N*_2_. When *N*_1_ has the higher metabolic rate, the species dynamics are similar and the coexistence conditions are identical to the model where both species have high metabolic rates (**A**). However, there are now two important differences. First, *N*_2_ (gray line) has a growth lag-phase. Second, both *R*_1_ and *R*_2_ are now depleted as the population abundance saturates. When *N*_2_ has the higher metabolic rates, the coexistence conditions are drastically different (**B**). *N*_1_ and *N*_2_ have growth lag-phases, and *R*_2_ and *R*_3_ are now the depleted resources. In addition, both species coexist at a final abundance determined only by the shared resource inflow rate (*ρ*_2_) across all resource inflow rates (*ρ*_1_*/ρ*_2_) tested. When we independently change *β*_1_ and *β*_2_, the final abundances depend on both values (**C**). As *β*_1_ decreases from 1 to 10^−4^ (blocks), the final abundance of *N*_1_ decreases and *N*_2_ increases for a given resource inflow ratio. In addition, as *B*_2_ increases from 10^−4^ to 1 (within each block), the final abundance of *N*_1_ decreases more and *N*_2_ increases for a given inflow ratio. Simulations based on Eqs. 14a–14e using the following parameters: *η*_*α*_ = 0.01, *C*_*αi*_ = 0.01, *µ*_*i*_ = 0.01, *λ*_*α*_ = 0.1 and *β*_*i*_ = 10 (high) or 10^−6^ (low), *ρ*_2_ = 80 or 100, and *ρ*_1_*/ρ*_2_ varies between 0.5 and 1.7 (**A**_*left*_ & **B**_*left*_:*ρ*_1_ = 64, *ρ*_2_ = 80). Resource concentrations are expressed in cell equivalents (i.e., moles required to generate a cell) and we assume 1:1 resource stoichiometry.

Finally, we consider the dynamics and equilibria when the internal metabolic rates, *β*_*i*_, are different. We find that, when the rates differ the outcome depends on which species has the higher metabolic rate. When the byproduct producer, *N*_1_, has the higher rate, then the results are similar to when both species have high internal metabolic rates (Fig. 5A). We do note, however, two important differences: 1) species *N*_2_ alone exhibits a growth lag-phase, and 2) both *R*_1_ and *R*_2_ are depleted as the species reach an equilibrium. However, when species *N*_2_ has the higher rate the coexistence conditions and high relative abundances for both species are greatly expanded. In fact, we find coexistence with moderate abundances along all inflow rates tested and the final abundances of both species are determined only by the shared resource, *R*_2_ (Fig. 5B). In addition, we find that both species exhibit growth lag-phases and that *R*_2_ and *R*_3_ are now the depleted resources. Between these limits, we find that the transition from classical resource limitation prediction to complete coexistence depends on both metabolic rates decreasing (Fig. 5C). These findings highlight how variation in both the internal metabolic reaction rates and in the environmental conditions can influence species interactions and change coexistence expectations.

## Discussion

Here, we reformulated the classic consumer-resource model by independently including resource uptake, internal metabolic rates, and byproduct export. We found that when internal metabolic dynamics are included – in addition to uptake – two common models of resource limitation (Liebig’s Law and multiplicative co-limitation) appear in particular limits of the internal reaction rates. In this model, we make predictions for the functional form of the production of metabolic byproducts which can be used in metabolic exchanges, and we balance the requirements for growth, energy, storage, and export. Because our model includes an independent term for internal metabolic rates, it provides a flexible and more general approach to understand how the resource landscape and other physical variables affect species dynamics in microbial communities. We further expand our model to include a second species which uses the metabolic byproduct of the first species. In this metabolically-informed approach, both species interactions (competition and facilitation) and resource use efficiency are emergent properties of the system. In addition, we show how internal metabolic reaction rates and the resource landscape determine species dynamics and equilibria. We find that the growth dynamics change when resources are *metabolically* limiting which depends on the internal metabolic rates. Therefore our model shows how metabolic rates and the resource landscape change the interactions between cross-feeding species which can shift the contributions of facilitation and competition at equilibrium. We further show that changes in the environmental landscape and/or metabolic rates can change resource and consumer dynamics in ways that alter species-species and species-resource correlations (see resource insets in Figs. 4 & 5); therefore, correlations may not accurately reflect interactions (see: Barner et al., 2018).

Classic formulations for pairwise interactions and consumer-resource dynamics have each led to insights regarding species coexistence, community stability, population self-regulation (Schoener, 1983; Barabás et al., 2017; Allesina and Tang, 2012; Leibold and McPeek, 2006). Lotka-Volterra (and related) equations provide a flexible approach to modeling a range of interactions between species but are unable to generalize across environmental variation because they do not provide an unambiguous way to include the resource landscape. While it may be possible to modify per capita growth rates to be a function of environmental conditions (e.g., temperature) and the resource landscape, how to incorporate these effects in the species interaction rates is ambiguous, at best. However, these models allow positive and negative species interactions to be explored straightforwardly, with far-reaching implications for coexistence and stability. Consumer-resource models explicitly include the interaction between the resource landscape and consumers, but at the expense of introducing more explicit mechanism, and therefore more choices in the way interactions are implemented (O’Dwyer, 2018). Including positive interactions through the production of resources has led to new predictions regarding the stability of communities (Butler and O’Dwyer, 2018, 2019), indicating that incorporating resource exchange (i.e., cross-feeding) may be important for understanding the dynamics of natural microbial communities. However, we do not know how sensitive these results are to the precise way consumption and exchange are formulated. Here, we incorporated metabolism more explicitly into consumer-resource dynamics and it allowed us to explain a broader range of community dynamics and natural phenomena, with less ambiguity in the functional form of interactions, and these metabolic rates may also reveal how environmental conditions that can alter metabolic rates (e.g., temperature) will contribute to species dynamics.

Our model used a simplified metabolic network to describe the internal physiology of cells and this allowed us to determine how uptake, growth, and export contribute to the dynamics and coexistence of interacting species. Specifically, we were able to incorporate competition via shared resource consumption and facilitation via the production and consumption of a metabolic byproduct (i.e., cross-feeding). Our model accomplishes this by using a consumer-resource framework with multiple inputs and outputs and by explicitly incorporating internal resource dynamics (see Figs. 1 & 3). Because we incorporate multiple resources as required inputs, there are similarities between our model and the synthesizing unit concept used in the dynamic energy budget theory (Kooijman, 1998, 2001). For example, both are able to produce limiting resource dynamics based on Liebig’s Law. However, our model was also able to produce multiplicative co-limitation, and it allowed us to explore metabolic cross-feeding by providing a mechanism for the production of cellular byproducts. We accomplish this by allowing internal resources, including luxury uptake and metabolic byproducts, to leak out of cells, which establishes strong conceptual similarities to Droop’s model (Droop, 1974). Because our model uses this internal flux balance approach, efficiency becomes an emergent property rather than a fixed term. This contrasts with approaches that include fixed efficiency terms that account for the internal processes without providing physiological mechanism (Moore et al., 1993; Schimel and Weintraub, 2003; Moore et al., 2005).

Likewise, our model differs from others that include explicit resource transfer pathways and corresponding efficiencies between species (Wiegert and Owen, 1971; DeAngelis et al., 1975; De Ruiter et al., 1995; Moore et al., 2005) because the byproducts and internal resource excesses are returned to the environment as ‘public goods’ rather than transferred directly. As such, our model uniquely allows us to incorporate a mixture of species interactions in addition to exploring how each population responds to a complex resource landscape because we include a simplified metabolic network.

Our model allows us to address scenarios that are unique to microbial communities – metabolic cross feeding and internal cellular metabolic rates – and this provided novel insights. First, by providing a mechanism for the production and subsequent consumption of metabolic byproducts (i.e., metabolic cross-feeding) our model provides a way to explore the role of facilitation interactions in a consumer-resource framework. Cross-feeding is a widely observed phenomenon in microbial communities (Morris et al., 2012; Morris, 2015; D’Souza et al., 2018). However, while cross-feeding appears to be ubiquitous, there are very few hypotheses or theoretical prediction for how it will shape the dynamics of interacting species. Our model provides a novel approach to explore this phenomenon and how it contributes to the diversity, complexity, and dynamics of microbial communities. In our model, the coexistence of species *N*_2_ is dependent on the production of a metabolic byproduct, *R*_3_, from species *N*_1_. Assuming there is no external input of the byproduct, the coexistence of *N*_2_ is dependent on *N*_1_. However, due to the way interactions are formulated, external inputs of byproduct alone are not enough to let *N*_2_ outcompete *N*_1_ (see Supplemental). As such, our model provides a new tool to explore the benefits and consequences of cross-feeding in microbial communities. Second, by separating the uptake and internal metabolic rates, our model is able to produce unique species dynamics and it recapitulates two common models of resource limitation (Liebig’s Law and multiplicative co-limitation). One major finding based on our model is that internal metabolic rates can have a strong impact on the dynamics of interacting species. Specifically, changes in the internal metabolic rates can alter species coexistence (see Figs. 4 & 5). There are multiple mechanisms that could modify these metabolic rates, including temperature dependence, cofactors, and redox status (Price and Sowers, 2004; Glass and Orphan, 2012; Falkowski et al., 2016; Ramírez-Flandes et al., 2019; Russell and Cook, 1995). Therefore, our model demonstrates how environmental conditions can radically change the relative abundances of interacting species even when resource inflow rates are fixed. Together, these two components of the metabolically-informed consumer-resource model highlight the potential to address questions and scenarios that may be unique to microbial communities.

Our model provides a mechanistic approach to include diverse species interaction in a consumer resource framework and generates novel insights, but it also provides a solid foundation to explore new questions. For example, we have added mechanisms to describe both competition and facilitation based species interactions. These interactions are not set parameters as in Lotka-Volterra and therefore may be more realistic in terms of modeling natural systems. However, because we have proposed mechanisms for these interactions, some of our conclusions may not be robust due to the precise way we have implemented the specific interactions. This provides an opportunity to explore how changing the implementation yields additional insights and tests new hypotheses. Likewise, we assumed that uptake and export rates increase linearly with resource concentration, but these parts of our equations could easily be modified to incorporate Monod dynamics (as noted above). These simplifications provide a useful starting point from which other hypotheses can be generated and tested. For example, our simplified metabolic models may not capture important idiosyncrasies regarding how metabolism changes along environmental gradients. Perhaps a multi-species genome-scale metabolic model is needed to better describe the metabolic transformations and exchanges between species (Kauffman et al., 2003; Orth et al., 2010; Embree et al., 2015). However, we believe that our simplified version is more tractable, especially when complete genomic information is lacking, and similar approaches have been successfully implemented to describe natural systems (Kempes et al., 2012; Litchman et al., 2015; Chakraborty et al., 2017). The aim of our approach was to find a balance that allows us to more mechanistically describe species interactions while still being tractable for describing natural systems and explaining observed variation.

We propose that when extended more broadly, this approach will lead to more mechanistic predictions for the role of positive interactions along stress gradients (Callaway and Walker, 1997; Brooker and Callaghan, 1998; Koffel et al., 2018), the possibility of species interactions to stabilize or de-stabilize communities (Butler and O’Dwyer, 2018; Allesina and Tang, 2012), and the mechanisms underlying biodiversity-ecosystem function relationships (Duffy et al., 2007; Flynn et al., 2011; Cardinale et al., 2012). Species interactions often change across environmental gradients (Brooker and Callaghan, 1998; He et al., 2013) and a shift that promotes facilitation may increase ecosystem function (e.g., productivity; see: Cardinale et al., 2002). Likewise, increases in nutrient availability has been shown to favor competition over facilitation (Koffel et al., 2018) and can decrease the importance of mutualisms (Keller and Lau, 2018). Our model provides a mechanistic approach to understand these shifts based on the resource landscape and internal metabolic rates. In addition, our model demonstrates that even when competition is favored over facilitation, two species can still coexist. Other studies suggest that positive species interactions, such as facilitation mediated through cross-feeding, can promote coexistence via negative frequency dependence (Morris, 2015). Our model demonstrates how even when competition is favored between cross-feeding species, the poorer competitor can remain as a rare member of the community due to the leakiness of the cross-feeder interaction; although, we predict that stochastic fluctuations could lead to exclusion. As such, our model can be used to better understand why some systems favor facilitation versus competition and how facilitation promotes coexistence and potentially the coevolution between species. In short, we propose that metabolically-informed consumer-resource dynamics will provide a platform to explore the consequences of cooperative and competitive interactions across environmental contexts.

## Acknowledgements

We thank Jake McKinlay, Thomas Koffel, and two anonymous reviewers for critical feedback on an earlier version of this manuscript. This work was supported by the Simons Foundation Grant # 376199 and the James S. McDonnell Foundation Grant # 220020439.

## References

Abrams PA (1982) Functional Responses of Optimal Foragers. The American Naturalist 120(3):382–390, DOI 10.1086/283996

Abrams PA (1983) Arguments in Favor of Higher Order Interactions. The American Naturalist 121(6):887–891, DOI 10.1086/284111

Abrams PA (2009) Determining the Functional Form of Density Dependence: Deductive Approaches for Consumer-Resource Systems Having a Single Resource. The American Naturalist 174(3):321–330, DOI 10.1086/603627

Allesina S, Tang S (2012) Stability criteria for complex ecosystems. Nature 483(7388):205–8, DOI 10.1038/nature10832

Barabás G, Michalska-Smith MJ, Allesina S (2017) Self-regulation and the stability of large ecological networks. Nature Ecology & Evolution 1(12):1870–1875, DOI 10.1038/s41559-017-0357-6

Barner AK, Coblentz KE, Hacker SD, Menge BA (2018) Fundamental contradictions among observational and experimental estimates of non-trophic species interactions. Ecology 99(3):557–566, DOI 10.1002/ecy.2133

Brooker RW, Callaghan TV (1998) The balance between positive and negative plant interactions and its relationship to environmental gradients: a model. Oikos 81(1):196, DOI 10.2307/3546481

Butler S, O’Dwyer JP (2018) Stability criteria for complex microbial communities. Nature Communications 9(1):2970, DOI 10.1038/s41467-018-05308-z

Butler S, O’Dwyer JP (2019) Cooperation and Stability for Complex Systems in Resource Limited Environments. Theoretical Ecology In Press, DOI xxx

Callaway RM, Walker LR (1997) Competition and facilitation: A synthetic approach to interactions in plant communities. Ecology 78(7):1958–1965, DOI 10.1890/0012-9658(1997)078[1958:CAFASA]2.0.CO;2

Cardinale BJ, Palmer MA, Collins SL (2002) Species diversity enhances ecosystem functioning through interspecific facilitation. Nature 415(6870):426–9, DOI 10.1038/415426a

Cardinale BJ, Duffy JE, Gonzalez A, Hooper DU, Perrings C, Venail P, Narwani A, MacE GM, Tilman D, Wardle DA, Kinzig AP, Daily GC, Loreau M, Grace JB, Larigauderie A, Srivastava DS, Naeem S (2012) Biodiversity loss and its impact on humanity. Nature 486(7401):59–67, DOI 10.1038/nature11148, 9605103

Carini P, Campbell EO, Morré J and Sañudo-Wilhelmy SAxs, Thrash JC, Bennett SE, Temperton B, Begley T, Giovannoni SJ (2014) Discovery of a SAR11 growth requirement for thiamin’s pyrimidine precursor and its distribution in the Sargasso Sea. ISME J 8(8):1727–1738, DOI 10.1038/ismej.2014.61

Carrara F, Giometto A, Seymour M, Rinaldo A, Altermatt F (2015) Inferring species interactions in ecological communities: A comparison of methods at different levels of complexity. Methods in Ecology and Evolution 6(8):895–906, DOI 10.1111/2041-210X.12363

Chakraborty S, Nielsen LT, Andersen KH (2017) Trophic Strategies of Unicellular Plankton. The American Naturalist 189(4):E77–E90, DOI 10.1086/690764

Cherif M, Loreau M (2007) Stoichiometric constraints on resource use, competitive interactions, and elemental cycling in microbial decomposers. The American Naturalist 169(6):709–724, DOI 10.1086/516844

Cherif M, Loreau M (2010) Towards a more biologically realistic use of Droop’s equations to model growth under multiple nutrient limitation. Oikos 119(6):897–907, DOI 10.1111/j.1600-0706.2010.18397.x

De Ruiter PC, Neutel AM, Moore JC (1995) Energetics, patterns of interaction strengths, and stability in real ecosystems. Science (80-) 269(5228):1257–1260, DOI 10.1126/science.269.5228.1257

DeAngelis DL, Goldstein RA, O’Neill RV (1975) A Model for Tropic Interaction. Ecology 56(4):881–892, DOI 10.2307/1936298

DeAngelis DL, Mulholland PJ, Palumbo AV, Steinman AD, Huston MA, Elwood JW (1989) Nutrient Dynamics and Food-Web Stability. Annual Review of Ecology and Systematics 20:71–95, DOI 10.1146/annurev.es.20.110189.000443

Droop MR (1974) The nutrient status of algal cells in continuous culture. Journal of the Marine Biological Association of the United Kingdom 54(04):825, DOI 10.1017/S002531540005760X

D’Souza G, Shitut S, Preussger D, Yousif G, Waschina S, Kost C (2018) Ecology and evolution of metabolic cross-feeding interactions in bacteria. Nat Prod Rep 35(5):455–488, DOI 10.1039/c8np00009c

Duffy JE, Cardinale BJ, France KE, McIntyre PB, Thébault E, Loreau M (2007) The functional role of biodiversity in ecosystems: incorporating trophic complexity. Ecology Letters 10(6):522–38, DOI 10.1111/j.1461-0248.2007.01037.x

Duncan SH, Louis P, Flint HJ (2004) Lactate-Utilizing Bacteria, Isolated from Human Feces, That Produce Butyrate as a Major Fermentation Product. Applied and Environmental Microbiology 70(10):5810–5817, DOI 10.1128/AEM.70.10.5810-5817.2004

Embree M, Liu JK, Al-Bassam MM, Zengler K (2015) Networks of energetic and metabolic interactions define dynamics in microbial communities. Proceedings of the National Academy of Sciences 112(50):15450–15455, DOI 10.1073/pnas.1506034112, 1011.1669v3

Falkowski PG, Jelen B, Giovannelli D (2016) The Role of Microbial Electron Transfer in the Coevolution of the Geosphere and Biosphere. Annu Rev Microbiol 70(1), DOI DOI:10.1146/annurev-micro-102215-095521, URL http://www.annualreviews.org/doi/abs/10.1146/annurev-micro-102215-095521

Flynn DFB, Mirotchnick N, Jain M, Palmer MI, Naeem S (2011) Functional and phylogenetic diversity as predictors of biodiversity–ecosystem-function relationships. Ecology 92(8):1573–1581, DOI 10.1890/10-1245.1, 1011.1669v3

Freilich S, Zarecki R, Eilam O, Segal ES, Henry CS, Kupiec M, Gophna U, Sharan R, Ruppin E (2011) Competitive and cooperative metabolic interactions in bacterial communities. Nature Communications 2:589, DOI 10.1038/ncomms1597

Gause GF, Witt AA (1935) Behavior of Mixed Populations and the Problem of Natural Selection. The American Naturalist 69(725):596–609, DOI 10.1086/280628

Glass JB, Orphan VJ (2012) Trace Metal Requirements for Microbial Enzymes Involved in the Production and Consumption of Methane and Nitrous Oxide. Front Microbiol 3(FEB):1–20, DOI 10.3389/fmicb.2012.00061, URL http://journal.frontiersin.org/article/10.3389/fmicb.2012.00061/abstract

Gottschalk G (1986) Bacterial Metabolism, 2nd edn. Springer-Verlag, New York, NY

Grilli J, Barabás G, Michalska-Smith MJ, Allesina S (2017) Higher-order interactions stabilize dynamics in competitive network models. Nature 548(DOI: 10.1038/nature23273):210–213, DOI 10.1038/nature23273

Grover JP (1990) Resource Competition in a Variable Environment: Phytoplankton Growing According to Monod’s Model. The American Naturalist 136(6):771–789, DOI 10.1086/285131

Grover JP (2011) Resource storage and competition with spatial and temporal variation in resource availability. The American naturalist 178(5):E124–48, DOI 10.1086/662163

Hall SR (2009) Stoichiometrically Explicit Food Webs: Feedbacks between Resource Supply, Elemental Constraints, and Species Diversity. Annual Review of Ecology, Evolution, and Systematics 40(1):503–528, DOI 10.1146/annurev.ecolsys.39.110707.173518

Harpole WS, Ngai JT, Cleland EE, Seabloom EW, Borer ET, Bracken ME, Elser JJ, Gruner DS, Hillebrand H, Shurin JB, Smith JE (2011) Nutrient co-limitation of primary producer communities. Ecology Letters 14(9):852–862, DOI 10.1111/j.1461-0248.2011.01651.x

He Q, Bertness MD, Altieri AH (2013) Global shifts towards positive species interactions with increasing environmental stress. Ecology Letters 16(5):695–706, DOI 10.1111/ele.12080, 2666

Holling CS (1959) The Components of Predation as Revealed by a Study of Small-Mammal Predation of the European Pine Sawfly. The Canadian Entomologist 91(05):293–320, DOI 10.4039/Ent91293-5

Holt RD (1977) Predation, Apparent Competition, and the Structure of Prey Communities. Theoretical Population Biology 12:197–229, DOI 10.1016/0040-5809(77)90042-9

Ives A, Dennis B, Cottingham K, Carpenter S (2003) Estimating community stability and ecological interactions from time-series data. Ecological Monographs 73(2):301–330, DOI 10.1890/0012-9615(2003)073[0301:ECSAEI]2.0.CO;2

Jiao Y, Navid A, Stewart BJ, McKinlay JB, Thelen MP, Pett-Ridge J (2012) Syntrophic metabolism of a co-culture containing Clostridium cellulolyticum and Rhodopseudomonas palustris for hydrogen production. International Journal of Hydrogen Energy 37(16):11719–11726, DOI 10.1016/j.ijhydene.2012.05.100

Kauffman KJ, Prakash P, Edwards JS (2003) Advances in flux balance analysis. Current Opinion in Biotechnology 14(5):491–496, DOI 10.1016/j.copbio.2003.08.001,/www.nature.com/nature/journal/v473/n7346/abs/10.1038-nature10011-unlocked.html#supplementary-information

Keller KR, Lau JA (2018) When mutualisms matter: Rhizobia effects on plant communities depend on host plant population and soil nitrogen availability. Journal of Ecology 106(3):1046–1056, DOI 10.1111/1365-2745.12938

Kempes CP, Dutkiewicz S, Follows MJ (2012) Growth, metabolic partitioning, and the size of microorganisms. Proceedings of the National Academy of Sciences 109(2):495–500, DOI 10.1073/pnas.1115585109

Koffel T, Boudsocq S, Loeuille N, Daufresne T (2018) Facilitation- vs. competition-driven succession: the key role of resource-ratio. Ecology Letters 21(7):1010–1021, DOI 10.1111/ele.12966

Kooijman SA (1998) The Synthesizing Unit as model for the stoichiometric fusion and branching of metabolic fluxes. Biophys Chem 73(1-2):179–188, DOI 10.1016/S0301-4622(98)00162-8

Kooijman SA (2001) Quantitative aspects of metabolic organization: A discussion of concepts. Philosophical Transactions of the Royal Society B: Biological Sciences 356(1407):331–349, DOI 10.1098/rstb.2000.0771

Leibold M, McPeek M (2006) Coexistence of the niche and neutral perspectives in community ecology. Ecology 87(6):1399–1410, DOI 10.1890/0012-9658(2006)87[1399:COTNAN]2.0.CO;2

von Liebig JF, Gregory W (1842) Animal chemistry: or, Organic chemistry in its application to physiology and pathology. John Owen

Litchman E (2003) Competition and coexistence of phytoplankton under fluctuating light: experiments with two cyanobacteria. Aquatic Microbial Ecology 31:241–248, DOI 10.3354/ame031241

Litchman E, Edwards KF, Klausmeier CA (2015) Microbial resource utilization traits and trade-offs: implications for community structure, functioning, and biogeochemical impacts at present and in the future. Frontiers in Microbiology 06(April):1–10, DOI 10.3389/fmicb.2015.00254

Loreau M (2001) Microbial diversity, producer-decomposer interactions and ecosystem processes: a theoretical model. Proceedings of the Royal Society 268(1464):303–9, DOI 10.1098/rspb.2000.1366

Loreau M (2010) From Populations to Ecosystems: Theoretical Foundations for a New Ecological Synthesis. Princeton University Press, Princeton, NJ

Lotka AJ (1932) The growth of mixed populations: Two species competing for a common food supply. Journal of the Washington Academy of Sciences 22(16/17):461–469, DOI 10.1007/978-3-642-50151-7_12

MacArthur R (1970) Species packing and competitive equilibrium for many species. Theoretical Population Biology 1(1):1–11, DOI 10.1016/0040-5809(70)90039-0

Marino S, Baxter NT, Huffnagle GB, Petrosino JF, Schloss PD (2014) Mathematical modeling of primary succession of murine intestinal microbiota. Proceedings of the National Academy of Sciences 111(1):439–444, DOI 10.1073/pnas.1311322111, 1011.1669v3

Mee MT, Collins JJ, Church GM, Wang HH (2014) Syntrophic exchange in synthetic microbial communities. Proceedings of the National Academy of Sciences of the United States of America 111(20):E2149–56, DOI 10.1073/pnas.1405641111

Moore JC, Ruiter PCD, Hunt HW (1993) Influence of Productivity on the Stability of Real and Model Ecosystems. Science (80-) 261(5123):906–908, URL http://www.jstor.org/stable/2882122

Moore JC, McCann K, De Ruiter PC (2005) Modeling trophic pathways, nutrient cycling, and dynamic stability in soils. Pedobiologia (Jena) 49(6):499–510, DOI 10.1016/j.pedobi.2005.05.008

Morris JJ (2015) Black Queen evolution: The role of leakiness in structuring microbial communities. Trends in Genetics 31(8):475–482, DOI 10.1016/j.tig.2015.05.004

Morris JJ, Lenski RE, Zinser ER (2012) The black queen hypothesis: evolution of dependencies through adaptive gene loss. MBio 3(2):e00036.#x2013;12, DOI 10.1128/mBio.00036-12

Mougi A, Kondoh M (2012) Diversity of Interaction Types and Ecological Community Stability. Science 337(6092):349–351, DOI 10.1126/science.1220529

Murdoch WW, Briggs CJ, Nisbet RM (2003) Consumer-resource dynamics, vol 36. Princeton University Press

Nisbet RM, Muller EB, Lika K, Kooijman SaLM (2008) From molecules to ecosystems through dynamic energy budget models. Journal of Animal Ecology 69(6):913–926, DOI 10.1111/j.1365-2656.2000.00448.x

Odum EP (1959) Fundamentals of ecology. WB Saunders company

O’Dwyer JP (2018) Whence Lotka-Volterra? Theoretical Ecology pp 1–12, DOI 10.1007/s12080-018-0377-0

Orth JD, Thiele I, Palsson BO (2010) What is flux balance analysis? Nature Biotechnology 28(3):245–248, DOI 10.1038/nbt.1614, NIHMS150003

Pacheco AR, Moel M, Segrè D (2019) Costless metabolic secretions as drivers of interspecies interactions in microbial ecosystems. Nature Communications 10(1):103, DOI 10.1038/s41467-018-07946-9

Pande S, Merker H, Bohl K, Reichelt M, Schuster S, de Figueiredo LF, Kaleta C, Kost C (2014) Fitness and stability of obligate cross-feeding interactions that emerge upon gene loss in bacteria. ISME J 8:953–62, DOI 10.1038/ismej.2013.211, URL http://www.ncbi.nlm.nih.gov/pubmed/24285359

Pfeiffer T, Bonhoeffer S (2004) Evolution of Cross-Feeding in Microbial Populations. The American Naturalist 163(6):E126–E135, DOI 10.1086/383593

Price PB, Sowers T (2004) Temperature dependence of metabolic rates for microbial growth, maintenance, and survival. Proceedings of the National Academy of Sciences 101(13):4631–4636, DOI 10.1073/pnas.0400522101

Ramírez-Flandes S, González B, Ulloa O (2019) Redox traits characterize the organization of global microbial communities. Proc Natl Acad Sci 116(9):3630–3635, DOI 10.1073/pnas.1817554116

Russell JB, Cook GM (1995) Energetics of Bacterial Growth: Balance of Anabolic and Catabolic Reactions. Microbiology and Molecular Biology Reviews 59(1):48–62

Schimel JP, Weintraub MN (2003) The implications of exoenzyme activity on microbial carbon and nitrogen limitation in soil: A theoretical model. Soil Biol Biochem 35(4):549–563, DOI 10.1016/S0038-0717(03)00015-4

Schoener TW (1983) Field Experiments on Interspecific Competition. The American Naturalist 122(2):240–285, DOI 10.1086/284133

Sterner RW, Elser JJ (2002) Ecological stoichiometry: the biology of elements from molecules to the biosphere. Princeton University Press, Princeton, NJ

Sun Z, Koffel T, Stump SM, Grimaud GM, Klausmeier CA (2019) Microbial cross-feeding promotes multiple stable states and species coexistence, but also susceptibility to cheaters. Journal of Theoretical Biology 465:63–77, DOI 10.1016/j.jtbi.2019.01.009

Tasoff J, Mee MT, Wang HH (2015) An economic framework of microbial trade. PLoS ONE 10(7):1–20, DOI 10.1371/journal.pone.0132907

Terry JCD, Morris RJ, Bonsall MB (2017) Trophic interaction modifications: an empirical and theoretical framework. Ecology Letters 20(10):1219–1230, DOI 10.1111/ele.12824

Tilman D (1977) Resource competition between plankton algae: an experimental and theoretical approach. Ecology 58(2):338–348

Tilman D (1980) Resources: a graphical-mechanistic approach to competition and predation. The American Naturalist 116(3):362–393

Tilman D (1987) The Importance of the Mechanisms of Interspecific Competition. The American Naturalist 129(5):769–774, DOI 10.1086/284672

Tilman D, Kilham SS, Kilham P (1982) Phytoplankton community ecology: the role of limiting nutrients. Annual Review of Ecology and Systematics 13(1):349–372, DOI 10.1146/annurev.es.13.110182.002025

Vellend M (2010) Conceptual synthesis in community ecology. The Quarterly review of biology 85(2):183–206, DOI 10.1086/652373

Vellend M (2016) The theory of ecological communities. Princeton University Press

Volterra V (1926) Fluctuations in the Abundance of a Species considered Mathematically. Nature 118(2972):558–560, DOI 10.1038/118558a0

Wiegert RG, Owen DF (1971) Trophic structure, available resources and population density in terrestrial vs. aquatic ecosystems. J Theor Biol 30(1):69–81, DOI 10.1016/0022-5193(71)90037-3

Xiao Y, Angulo MT, Friedman J, Waldor MK, Weiss ST, Liu YY (2017) Mapping the ecological networks of microbial communities. Nature Communications 8(1):2042, DOI 10.1038/s41467-017-02090-2

Zelezniak A, Andrejev S, Ponomarova O, Mende DR, Bork P, Patil KR (2015) Metabolic dependencies drive species cooccurrence in diverse microbial communities. Proceedings of the National Academy of Sciences 112(20):6449–6454, DOI 10.1073/pnas.1421834112, 1408.1149

Zomorrodi AR, Segrè D (2016) Synthetic Ecology of Microbes: Mathematical Models and Applications. Journal of Molecular Biology 428(5):837–861, DOI 10.1016/j.jmb.2015.10.019

