## Supplemental for "Species dynamics and interactions via metabolically informed consumer-resource models"

**Mario E. Muscarella · James P. O'Dwyer**

the date of receipt and acceptance should be inserted later

### Lotka-Volterra Equations

The Lotka-Volterra equations provide an example of mathematical models which have been widely used for close to a century (Lotka, 1932; Volterra, 1926). These equations characterize species interactions in terms of the net, direct effect of one population on another's growth rate, so that in the case of two species with abundances $N_1$ and $N_2$:

$$\frac{dN_1}{dt} = r_1 N_1 - a_{11} N_1^2 - a_{21} N_1 N_2$$

$$\frac{dN_2}{dt} = r_2 N_2 - a_{22} N_2^2 - a_{12} N_1 N_2. \tag{1}$$

Here, $r_1$ and $r_2$ are per capita growth rates when species are rare, and the parameters $a_{ij}$ (collectively called a community matrix) represent intra- and interspecific interactions. Empirically, it has been extremely difficult to reliably estimate these parameters (Schoener, 1983; Tilman, 1987). Even where it has been possible to infer or approximate pairwise interactions (Stein et al., 2013; Marino et al., 2014; Fisher and Mehta, 2014; Bucci et al., 2016), it may be difficult to translate the inferred interactions in different environmental contexts. In part, this is difficult because these models lack an explicit description of the mechanisms mediating interactions (Abrams, 1983; Grilli et al., 2017). For example, if two species compete, it is often because they consume common resources (Gause and Witt, 1935; MacArthur, 1970; Schoener, 1983). However, these models assume that resource dynamics can be safely ignored because resource dynamics are faster than consumer dynamics (MacArthur, 1970). This exposes an important context-dependence of Lotka-Volterra type equations: the strength and even the sign of a pairwise interaction may depend on what resources are present (Xiao et al., 2017). As such, variation in the environmental landscape can influence species composition due to differences in competitive ability and the context dependence of species interactions (Cadotte and Tucker, 2017).

Address(es) of author(s) should be given

### Consumer-Resource Equations

An alternate approach is to model competitive interactions as the explicit result of shared resource consumption (Tilman et al., 1982; Grover, 1990; Litchman, 2003; Abrams, 2009). For example, in the case of two species with abundances $N_1$ and $N_2$ competing for a single shared resource, $R$, the prototypical consumer-resource model is:

$$\begin{aligned}\frac{dR}{dt} &= \rho - \eta R - \sum_i a_i N_i R \\ \frac{dN_1}{dt} &= \varepsilon_1 a_1 N_1 R - \mu_1 N_1 \\ \frac{dN_2}{dt} &= \varepsilon_2 a_2 N_2 R - \mu_2 N_2, \end{aligned} \tag{2}$$

where $\rho$ and $\eta$ describe the environmental inflow and outflow rates of an abiotic resource, $a_i$ describes the resource uptake rates, $\varepsilon_i$ describes the resource use efficiency, and $\mu_i$ are the species mortality rates. These models produce species interactions as an emergent property dependent on shared resource consumption, and so the issue of inferring species interactions is no longer quite the right question—though there is now a challenge in determining consumer feeding preferences. Assuming we can infer or otherwise estimate those preferences, one critical aspect of the environmental context is now explicitly characterized, via resource input rates like $\rho$. As such, species dynamics across resource landscapes can be understood better than in the case of Lotka-Volterra, where the effect of the environment is implicit (Tilman, 1977; Grover, 1990, 2011).

### Supplemental Figures

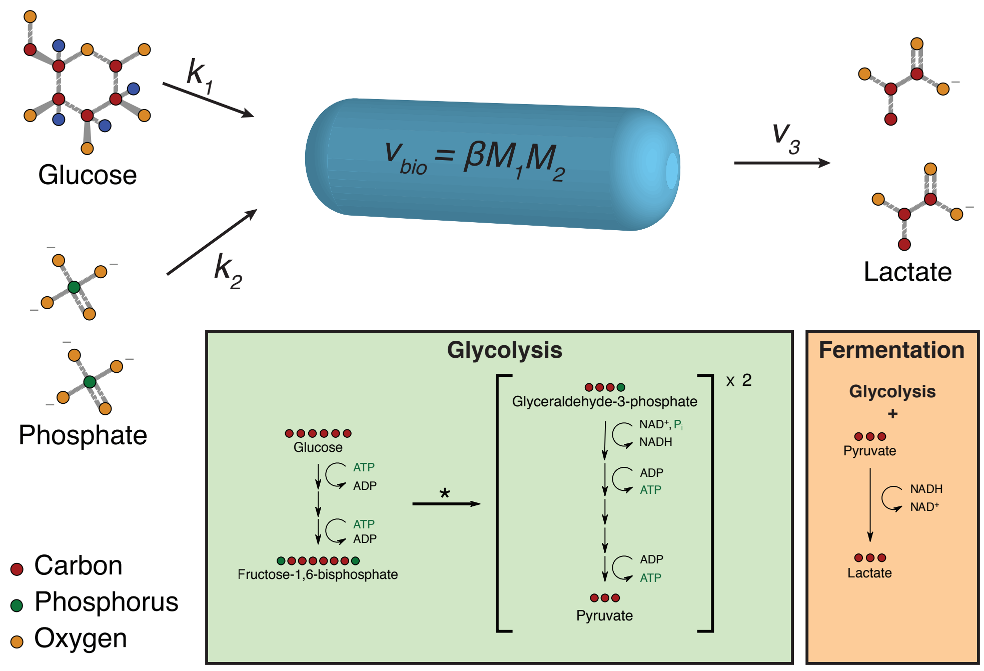


**Fig. 1 Conceptual Model**. To demonstrate our metabolically informed consumer-resource model we used fermentation as a prototype. Fermentation is an anaerobic—usually sugar consuming—metabolic lifestyle, and is the primary anaerobic energy-producing reaction for many microorganisms (Gottschalk, 1986). A signature of fermentation is that it results in byproducts such as organic acids, alcohols, and/or gases, which are produced due to the incomplete resource oxidation during the energy producing reactions. When organic acids (e.g., lactate) are produced, they are often used as resources by other microorganisms. Homolactic fermentation, the simplest type of fermentation, results in the incomplete oxidation of glucose. The inputs are one glucose and two phosphate molecules, and the products are two lactate molecules. This energy producing reaction yields a net two ATP per glucose. Here, we show embedded the detailed chemical reactions involved with homolactic fermentation. First, glycolysis is used to turn one molecule of glucose into two pyruvate molecules. At this stage, the pyruvate can either be used for biomass, or it can be fermented into lactate. In our model, we simplify this picture by assuming that glucose and phosphate enter the cell at uptake rates $k_i$, biomass is generated at rate $\beta M_1 M_2$, and the metabolic byproduct, lactate, is exported from the cell at rate $v_3$.

#### $R_3$ Influx Rate

To test how external inputs of $R_3$ change or modify the dynamics of the two consumer species ($N_1$ and $N_2$), we performed a series of simulations. First, using Eq. 14 we performed simulations where $R_3$ was added to the resource environment via external loading (i.e., $\rho_3 > 1$). The $R_3$ influx rate ($\rho_3$) varied between 0 and 120 cell equivalents. For our initial simulations, we used the following parameters: $\eta_\alpha = 0.01$, $C_{\alpha i} = 0.01$, $\mu_i = 0.01$, $\lambda_\alpha = 0.1$, $\beta_i = 1$, $\rho_2 = 100$, and $\rho_1 = 60$ or 100. We varied $R_2$ because it represents competition between $N_1$ and $N_2$. When $\rho_1$ is low (and therefore $R_1$ is limiting), the two species compete, and we predicted that $N_2$ would now have an advantage since it does not rely on $N_1$ alone for $R_3$.

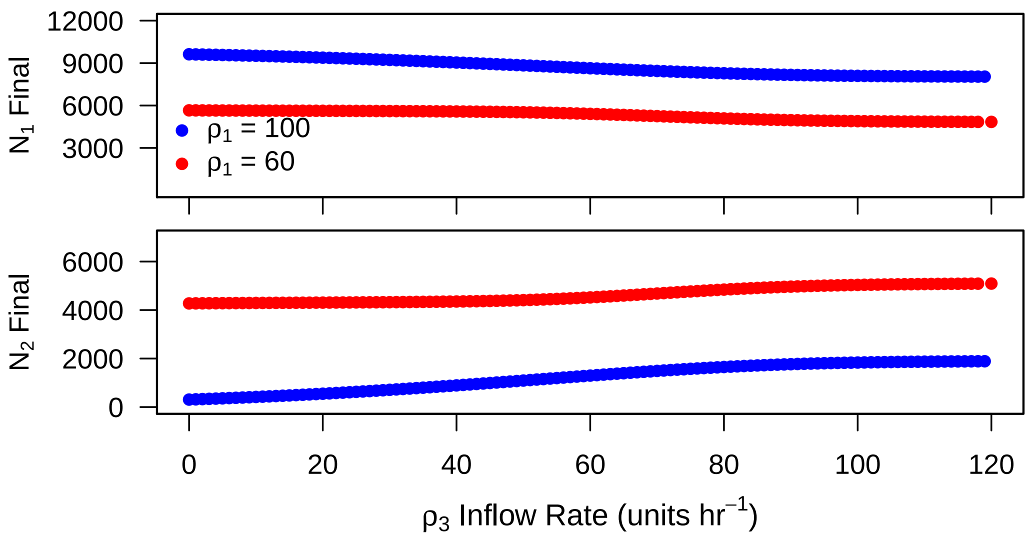


**Fig. 2 Influx experiment: same metabolic rates**. Competition with high metabolic rates when there is an external source of $R_3$. Final abundances for $N_1$ and $N_2$ are shown.

Based on these simulations, we found that the final abundance of $N_1$ and $N_2$ were affected by external loading of $R_3$ (Fig. S2). As predicted, the final abundance of $N_2$ increased with the influx rate of $R_3$, and there was a proportional decrease in the final abundance of $N_1$. However, the change in $N_1$ final abundance was small (even when $\rho_3$ was high). As such, $N_2$ was not able to out complete $N_1$ (i.e., no local extinction).

Next, we kept all of the parameters the same except we decreased the metabolic rate of $N_1$: $\beta_1 = 0.01$). We predicted that this would give $N_2$ a larger competitive advantage.

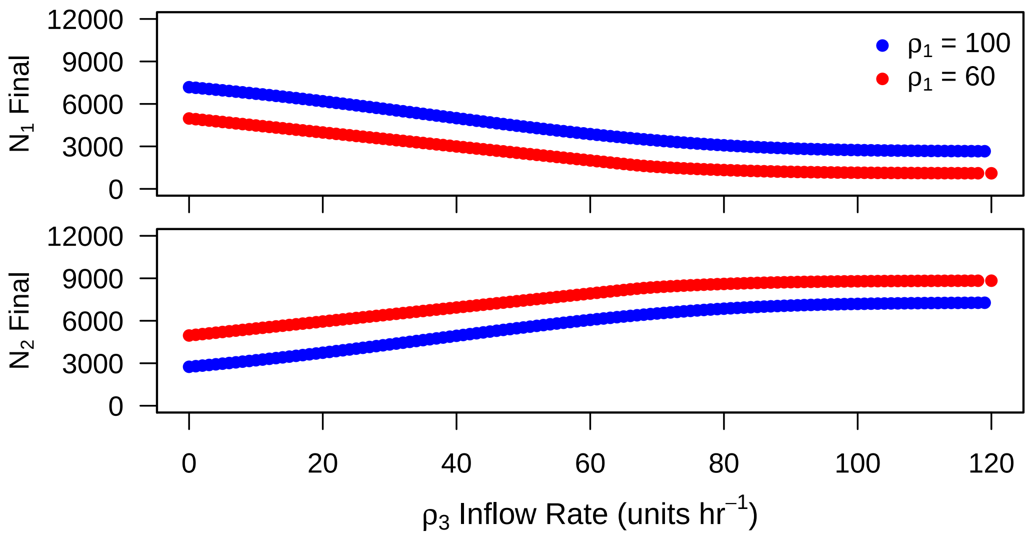


**Fig. 3 Influx experiment: different metabolic rates**. Competition when there is an external source of $R_3$ and $N_2$ has a higher metabolic rate. Final abundances for $N_1$ and $N_2$ are shown.

Based on these simulations, we found would that – as predicted – the competitive ability of $N_2$ could increase (Fig. S3). When $N_2$ has a higher metabolic rate than $N_1$, the increase in final abundance of $N_2$ is larger. However, while there is a substantial decrease in the final abundance of $N_1$, it did no go extinct. The *rare* occurrence of $N_1$ is due to the continued export of internal resources by $N_2$. It is possible that stochastic fluctuations could lead to the final extinction of $N_1$, but under the simulation conditions there is no extinction. These results demonstrate that it is possible for $N_2$ to have a competitive advantage, but only if the external inputs of $R_3$ is also accompanied by a metabolic advantage.

In conclusion, the external influx of the metabolic byproduct is not enough is allow $N_2$ to outcompete $N_1$. If order to outcompete, there would also need to be a large discrepancy in the internal metabolic rates.

### References


Abrams PA (1983) Arguments in Favor of Higher Order Interactions. The American Naturalist 121(6):887–891, DOI 10.1086/284111

Abrams PA (2009) Determining the Functional Form of Density Dependence: Deductive Approaches for Consumer-Resource Systems Having a Single Resource. The American Naturalist 174(3):321–330, DOI 10.1086/603627

Bucci V, Tzen B, Li N, Simmons M, Tanoue T, Bogart E, Deng L, Yeliseyev V, Delaney ML, Liu Q, et al. (2016) Mdsine: Microbial dynamical systems inference engine for microbiome time-series analyses. Genome biology 17(1):121, DOI 110.1186/s13059-016-0980-6

Cadotte MW, Tucker CM (2017) Should Environmental Filtering be Abandoned? Trends in Ecology and Evolution 32(6):429–437, DOI 10.1016/j.tree.2017.03.004

Fisher CK, Mehta P (2014) Identifying keystone species in the human gut microbiome from metagenomic timeseries using sparse linear regression. PloS one 9(7):e102451, DOI 10.1371/journal.pone.0102451

Gause GF, Witt AA (1935) Behavior of Mixed Populations and the Problem of Natural Selection. The American Naturalist 69(725):596–609, DOI 10.1086/280628

Gottschalk G (1986) Bacterial Metabolism, 2nd edn. Springer-Verlag, New York, NY

Grilli J, Barabás G, Michalska-Smith MJ, Allesina S (2017) Higher-order interactions stabilize dynamics in competitive network models. Nature 548(DOI: 10.1038/nature23273):210–213, DOI 10.1038/nature23273

Grover JP (1990) Resource Competition in a Variable Environment: Phytoplankton Growing According to Monod's Model. The American Naturalist 136(6):771–789, DOI 10.1086/285131

Grover JP (2011) Resource storage and competition with spatial and temporal variation in resource availability. The American naturalist 178(5):E124–48, DOI 10.1086/662163

Litchman E (2003) Competition and coexistence of phytoplankton under fluctuating light: experiments with two cyanobacteria. Aquatic Microbial Ecology 31:241–248, DOI 10.3354/ame031241

Lotka AJ (1932) The growth of mixed populations: Two species competing for a common food supply. Journal of the Washington Academy of Sciences 22(16/17):461–469, DOI 10.1007/978-3-642-50151-7_12

MacArthur R (1970) Species packing and competitive equilibrium for many species. Theoretical Population Biology 1(1):1–11, DOI 10.1016/0040-5809(70)90039-0

Marino S, Baxter NT, Huffnagle GB, Petrosino JF, Schloss PD (2014) Mathematical modeling of primary succession of murine intestinal microbiota. Proceedings of the National Academy of Sciences 111(1):439–444, DOI

10.1073/pnas.1311322111, arXiv:1011.1669v3

Schoener TW (1983) Field Experiments on Interspecific Competition. The American Naturalist 122(2):240–285, DOI 10.1086/284133

Stein RR, Bucci V, Toussaint NC, Buffie CG, Rätsch G, Pamer EG, Sander C, Xavier JB (2013) Ecological modeling from time-series inference: insight into dynamics and stability of intestinal microbiota. PLoS computational biology 9(12):e1003388, DOI 10.1371/journal.pcbi.1003388

Tilman D (1977) Resource competition between plankton algae: an experimental and theoretical approach. Ecology 58(2):338–348

Tilman D (1987) The Importance of the Mechanisms of Interspecific Competition. The American Naturalist 129(5):769–774, DOI 10.1086/284672

Tilman D, Kilham SS, Kilham P (1982) Phytoplankton community ecology: the role of limiting nutrients. Annual Review of Ecology and Systematics 13(1):349–372, DOI 10.1146/annurev.es.13.110182.002025

Volterra V (1926) Fluctuations in the Abundance of a Species considered Mathematically. Nature 118(2972):558–560, DOI 10.1038/118558a0

Xiao Y, Angulo MT, Friedman J, Waldor MK, Weiss ST, Liu YY (2017) Mapping the ecological networks of microbial communities. Nature Communications 8(1):2042, DOI 10.1038/s41467-017-02090-2
